# PFKFB2 is Pivotal for Metabolic Flexibility and Differential Glucose Utilization

**DOI:** 10.1101/2025.05.26.656235

**Authors:** Kylene M. Harold, Satoshi Matsuzaki, Atul Pranay, Jie Zhu, Anna Faakye, Kenneth M. Humphries

**Author notes:** Corresponding Author: Kenneth M. Humphries, Ph.D. 825 N.E. 13th Street Oklahoma City, OK 73104.

## Abstract

**Background:** The heart’s constant energy demands make metabolic flexibility critical to its function as nutrient availability varies. The enzyme phosphofructokinase-2/fructose 2,6-bisphosphatase (PFKFB2) contributes to this flexibility by acting as a positive or negative regulator of cardiac glycolysis. We have previously shown that PFKFB2 is degraded in the diabetic heart and that a cardiac-specific PFKFB2 knockout (cKO) impacts ancillary glucose pathways and mitochondrial substrate preference. Therefore, defining PFKFB2’s role in mitochondrial metabolic flexibility is paramount to understanding both metabolic homeostasis and metabolic syndromes. Further, it is unknown how PFKFB2 loss impacts the heart’s response to acute stress. Here we examined how cardiac mitochondrial flexibility and the post-translational modification O-GlcNAcylation are affected in cKO mice in response to fasting or pharmacologic stimulation.

**Methods:** cKO and litter-matched controls (CON) were sacrificed in the fed or fasted (12 hours) states, with or without a 20 minute stimulant stress of caffeine and epinephrine.

Mitochondrial respiration, metabolomics, and changes to systemic glucose homeostasis were evaluated.

**Results:** cKO mice had moderate impairment in mitochondrial metabolic flexibility, affecting downstream glucose oxidation, respiration, and CPT1 activity. O-GlcNAcylation, a product of ancillary glucose metabolism, was upregulated in cKO hearts in the fed state, but this was ameliorated in the fasted state. Furthermore, metabolic remodeling in response to PFKFB2 loss was sufficient to impact circulating glucose in fasted and stressed states.

**Conclusions:** PFKFB2 is essential for fed-to-fasted changes in cardiac metabolism and plays an important regulatory role in protein O-GlcNAcylation. Its loss also affects systemic glucose homeostasis under stressed conditions.

**Graphic Abstract:** 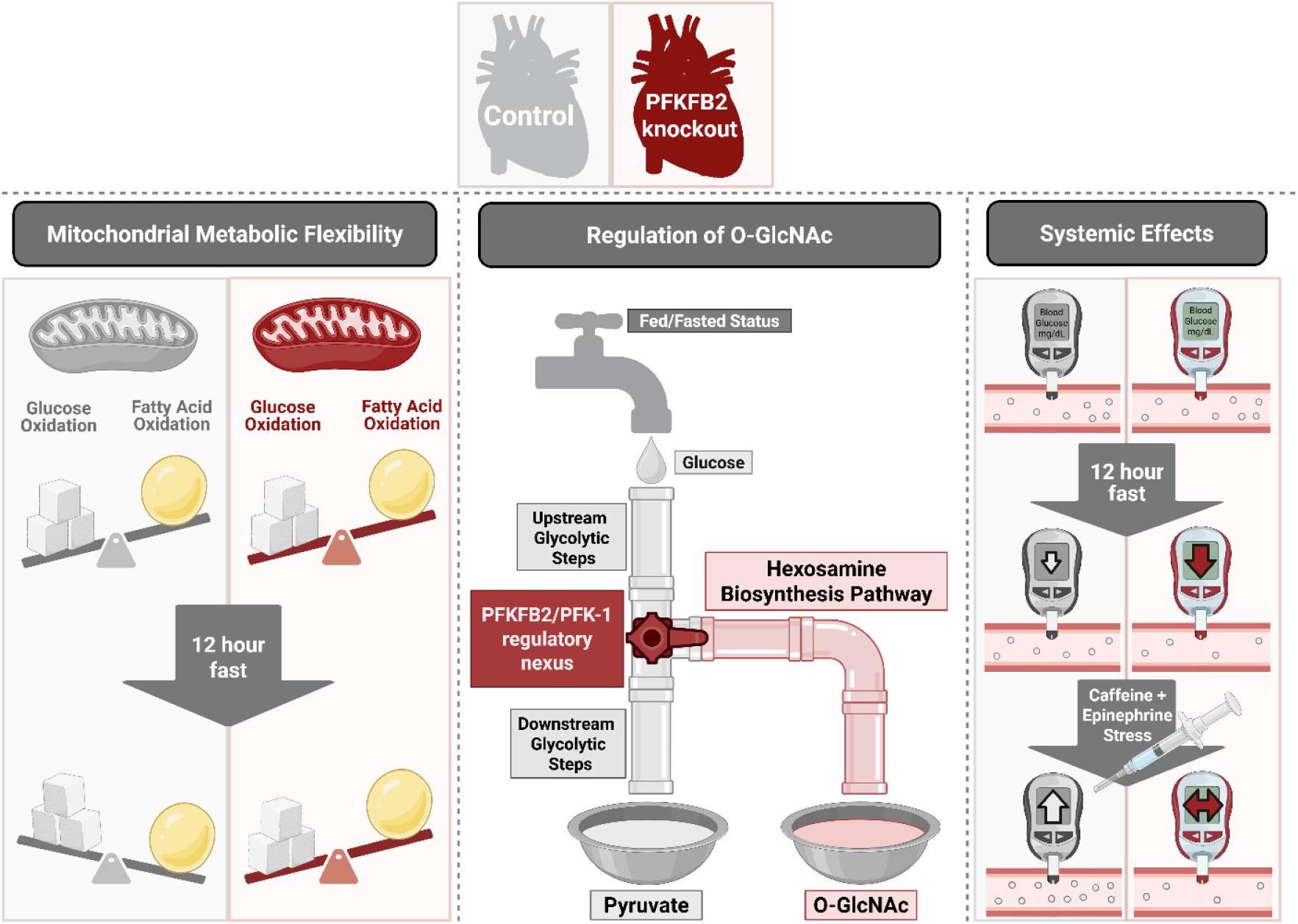

**Research Perspective:** This study raises and answers three key questions: how PFKFB2 contributes to cardiac mitochondrial metabolic flexibility, how post-prandial status regulates O-GlcNAcylation in a PFKFB2-dependent manner, and how altered cardiac glucose use impacts systemic glucose homeostasis under stress.

These findings highlight a novel role for nutrient state in regulating cardiac metabolism, and especially O-GlcNAcylation, with PFKFB2 loss.

Future studies should investigate whether reducing O-GlcNAcylation through fasting is sufficient to ameliorate pathological changes observed in the absence of PFKFB2.

## Introduction

The cardiac isoform of Phosphofructokinase-2/fructose 2,6-bisphosphatase (PFKFB2) serves as a switch in canonical glucose catabolism, whereby its phosphorylation in response to insulin or stress allows the heart to compensatorily upregulate glycolysis^1–3^. As a bifunctional glycolytic regulator, PFKFB2 both produces and degrades fructose 2,6-bisphosphate, an allosteric activator of Phosphofructokinase-1 which catalyzes the rate-limiting step of glycolysis^3–7^. Phosphorylation by AMPK, Akt, or PKA dictates the role of PFKFB2 as a kinase or phosphatase. When phosphorylated by these upstream kinases, PFKFB2 itself acts as a kinase, producing fructose 2,6-bisphosphate and activating glycolysis. Conversely, unphosphorylated PFKFB2 converts fructose 2,6-bisphosphate back to fructose 6-phosphate, decreasing activation of glycolysis. However, PFKFB2 is also degraded in the heart in response to decreased insulin signaling via multiple mechanisms and to varying degrees, scalable to the degree and duration of diminished insulin signaling^3,8,9^.

It is also established that decreased insulin signaling, as well as concomitant decreased glucose uptake and utilization, are key hallmarks of the fasted state in the heart^10^. This serves as a protective mechanism when carbohydrate stores become more finite, conserving glucose for obligate users such as erythrocytes. On a short-term timescale, PFKFB2 dephosphorylation and degradation likely contribute to the relationship between decreased insulin signaling and reduced glycolysis in the fasted heart. These fluctuations in PFKFB2 content and phosphorylation state contribute to necessary aspects of cardiac metabolic flexibility. However, on a longer term timescale, the chronic impairment of insulin signaling that marks metabolic syndrome and diabetes can promote sustained PFKFB2 loss^3^, which drives both metabolic and functional impairments in the heart^11^.

One possible contributor to the impacts of PFKFB2 loss beyond metabolic rigidity is activation of ancillary pathways such as the hexosamine biosynthesis pathway which promotes O-GlcNAcylation. This post-translational modification is upregulated in the diabetic heart and can promote pathology with chronic activation^12–14^. We have previously shown that in the fed state, loss of cardiac PFKFB2 drives changes suggestive of ancillary pathway activation including increased O-GlcNAc levels^11^. We therefore hypothesize that O-GlcNAcylation is driven by coordination between the phosphofructokinase regulatory nexus and nutrient status. We posited that maintenance of a baseline O-GlcNAc level is dependent on concomitant upregulation (post-prandially) or downregulation (with fasting) of both glucose uptake and PFKFB2.

Further, both PFKFB2 and O-GlcNAcylation have been implicated in the heart’s response to acute stress. We therefore also asked how cardiac O-GlcNAc levels and other aspects of glucose metabolism are impacted by an acute stimulant stress in the absence of PFKFB2.

Here, we used a cardiac-specific PFKFB2 knockout mouse model subjected to a 12-hour fast. This mild and acute metabolic stress allowed elucidation of PFKFB2 roles in metabolic flexibility, insulin signaling, and ancillary pathway activation. The study had three primary objectives. The first was to test the contribution of the phosphofructokinase nexus to mitochondrial metabolic flexibility with fasting. While the role of PFKFB2 in cardiac glycolytic metabolic flexibility is established, how PFKFB2 loss contributes to downstream remodeling of mitochondrial processes and flexibility in the mitochondria is unknown. The second objective was to determine the relative role of post-prandial status (substrate availability) for ancillary pathways in cardiac O-GlcNAc regulation when PFKFB2 is absent. Finally, we sought to determine the contribution of the cardiac phosphofructokinase nexus to glucose homeostasis in the fed and fasted states, with and without an acute pharmacologic stress of caffeine and epinephrine.

## Methods Animals

All mouse procedures were approved by the Oklahoma Medical Research Foundation Institutional Animal Care and Use Committee (IACUC). All mice were PFKFB2 flox/flox on a C57Bl6J background^11^. Mice were born in mendelian ratios, where 50% of mice per litter were heterozygous for MHC6-Cre and were thus cardiomyocyte-specific PFKFB2 knockout mice (cKO). The remaining 50% of mice were negative for Cre and served as litter-matched controls (CON). Mice were maintained on a standard stock chow diet and a 14/10 light/dark cycle. Lights are turned on at 0600 hours and turned off at 2000 hours. Mice were euthanized in either the fed or fasted state. Fed mice were euthanized at approximately 0700 hours, shortly after the end of the dark (active) cycle. Fasted mice were euthanized at approximately 1900 hours, close to the end of the light (inactive) cycle. Food and traditional bedding were removed from cages of fasted mice at 0700, 12 hours prior to euthanasia. Mice had access to water ad libitum. A combination of male and female mice was used with effort made to maintain equal proportions of each per group. Blood glucose was measured from the tail vein by a Contour Next glucometer at the time of euthanasia (as well as at 0700 hours in fasted mice) on the day of use.

In a subset of animals, to test the effects of acute stress on cardiac and systemic metabolism, we used intraperitoneal injections of caffeine (100 mg/kg body weight) and epinephrine (1.25 mg/kg body weight). To prevent distress in injected mice, this was done under inhaled isoflurane-induced anesthesia (2.0% for induction, 1-1.5% for maintenance). Stress was carried out for 20 minutes post-injection. Blood glucose was measured both prior to injection and at the end of this stress period.

After blood glucose measurement, euthanasia was performed by cervical dislocation under isoflurane-induced anesthesia, either via the drop method, or from gradual inhalation in the case of pharmacologically stressed mice. The only exception to this was animals used for respirometry, where cervical dislocation was performed in the absence of isoflurane due to the potential impacts of isoflurane on mitochondrial respiration^15,16^. Following euthanasia, blood was collected by cardiac puncture in some mice, hearts were perfused and excised, and hearts and other tissues were weighed and processed according to their intended uses.

## Mass Spectrometry-Based Metabolomic Assays

Targeted liquid chromatography- and semi-targeted gas chromatography-mass spectrometry assays (LC-MS and GC-MS, respectively) were performed as previously described^11^. Briefly, snap frozen hearts were pulverized and powder used for metabolite extraction with methanol and water (8:2, v:v, respectively). Adonitol was added as an internal standard, and metabolite extracts were divided into two aliquots, to be used for LC-MS and GC-MS assays. Liquid chromatographic separation for LC-MS analysis was performed using an Agilent InfinityLab Poroshell 120 HILIC-Z with the mobile phase consisting of an aqueous ammonium acetate buffer and an acetonitrile buffer. Total run time was 29 minutes. Targeted LC-MS analysis was performed on Agilent 6546 LC/Q-TOF-coupled to an Agilent 1290 Infinity II LC. For semi-targeted GC-MS analysis, samples were derivatized and analyzed on the Agilent 7890B-5977A GC–MS system in electron ionization mode. An added alkane mix was used as a ladder for retention time. All GC-MS and LC-MS data were analyzed by Agilent MassHunter quantitative analysis software (version 10.1 for LC-MS data and B.07.01 for GC-MS). GC-MS data were normalized to tissue weight and adonitol response. LC-MS data were normalized by total ion current and adonitol.

## Cardiac Homogenate Preparation and Fractionation

Except for hearts used for metabolomic assays, which were snap frozen and pulverized, hearts were immediately homogenized in 5 mL of mitochondrial isolation buffer (210 mM mannitol, 70 mM sucrose, 5 mM MOPS (3-(N-morpholino)propanesulfonic acid), 1 mM EDTA; pH 7.4) at the time of harvest. The resulting homogenate was centrifuged at 550×g for 5 minutes at 4°C. The pellet was saved as a crude nuclear fraction, and 200 μL of the supernatant was collected as our working homogenate. All remaining supernatant was centrifuged again at 10,000×g for 10 minutes at 4°C. This second pellet, a mitochondria-enriched fraction, was resuspended in 60 μL mitochondrial isolation buffer and the resulting suspension, to serve as our initial mitochondrial suspension for respirometry and mitochondrial enzyme activity assays, was quantified by bicinchoninic acid assay (BCA).

## Pyruvate Dehydrogenase Activity

Pyruvate dehydrogenase (PDH) activity was measured as previously described^11^. Briefly, isolated mitochondria were diluted 3:100 in OXPHOS buffer (210 mM mannitol, 70 mM sucrose, 10 mM MOPS, and 5 mM K_2_HPO_4_; pH 7.4). Diluted mitochondria were then incubated for 2 minutes with 0.25 mM pyruvate and 2.5 mM malate prior to further (1:5) dilution in a detergent solution composed of 25 mM MOPS with 0.05% Triton X-100. Following addition of 200 μM thiamine pyrophosphate, 2.5 mM pyruvate, 100 μM CoASH, 5.0 mM MgCl_2_, and 1 mM NAD^+^, NADH production was measured spectrophotometrically at 340nm every 7.5 seconds for 275 seconds. Initial absorbance prior to addition of NAD^+^, MgCl_2_, pyruvate, thiamine pyrophosphate, and CoASH was subtracted. The resulting signal was normalized to the starting protein concentration, which was measured via BCA.

## Respirometry

Oxygen consumption was monitored using a fluorescence lifetime-based sensor (Instech). A subset of isolated mitochondria was diluted to 0.25 mg/mL in OXPHOS buffer. Either 0.1 mM pyruvate and 1.0 mM malate, or 25 μM palmitoyl carnitine and 1.0 mM malate were added to mitochondria as substrates prior to their transfer to the respirometry chamber. After 2 minutes, 0.5 mM adenosine diphosphate was injected into the chamber to initiate State 3 respiration. The assay was completed when respiratory rates reached state 4.

## CPT1 activity

To measure carnitine palmitoyl transferase 1 (CPT1) activity, the initial suspension of mitochondria was further diluted in 25 mM MOPS buffer, such that final mitochondrial protein concentration was 1 mg/mL for all samples. 25 μL of this suspension (25 μg protein total), 100 μM palmitoyl CoA, and 100 μM 5,5-dithio-bis-(2-nitrobenzoic acid) were then preincubated for 20 minutes in a CPT assay buffer (25 mM MOPS at pH 7.4, 1 mM EGTA, 0.1% BSA). After acquiring a baseline measurement, 10 mM carnitine was added and absorbance at 412 nm was monitored for 10 minutes. The linear range, which occurred approximately between seconds 60 and 240 of measurement, was used for quantification.

## Western Blot and Immunoprecipitation

For western blot assays, either working homogenate or mitochondrial fraction samples were prepared with lithium dodecyl sulfate, a protease/phosphatase inhibitor cocktail (Halt, Thermo Fisher Scientific), 18 mM dithiothreitol, and in the case of mitochondrial fractions, diluted to final concentrations with water. These samples were then heated at 95 °C for 10 minutes and run on 4-12% Bis-Tris Gels at 200 V for either 45 or 65 minutes, depending on the size of the protein of interest and necessary separation. A semi-dry transfer of proteins onto nitrocellulose was then performed, using 30 V for 1 hour. Immediately after transfer, Revert (LI-COR) total protein stain was used for later normalization purposes. Primary antibodies were suspended in Intercept blocking buffer (LI-COR) at concentrations shown in Table 1. Blots were incubated in primary antibody (Table 1) overnight at 4°C with rocking. Blots were then washed 3 times for 10 minutes each at room temperature in tris-buffered saline (TBS) with 1% Tween 20, prior to incubation for one hour at room temperature in their respective fluorophore-tagged anti-rabbit (IRDye 800CW, LI-COR) or anti-mouse (IRDye 680RD, LI-COR) secondary antibody (concentrations in Table 1, suspended in Intercept blocking buffer). Lastly, blots were washed 2 times for 10 minutes each in TBS with 1% Tween 20, followed by a single 5 minute wash in TBS without detergent. Fluorescent signal from secondary antibodies was acquired by the Odyssey CLx system (LI-COR) and quantified via Image Studio (Version 5.2, LI-COR).

**Table 1.**
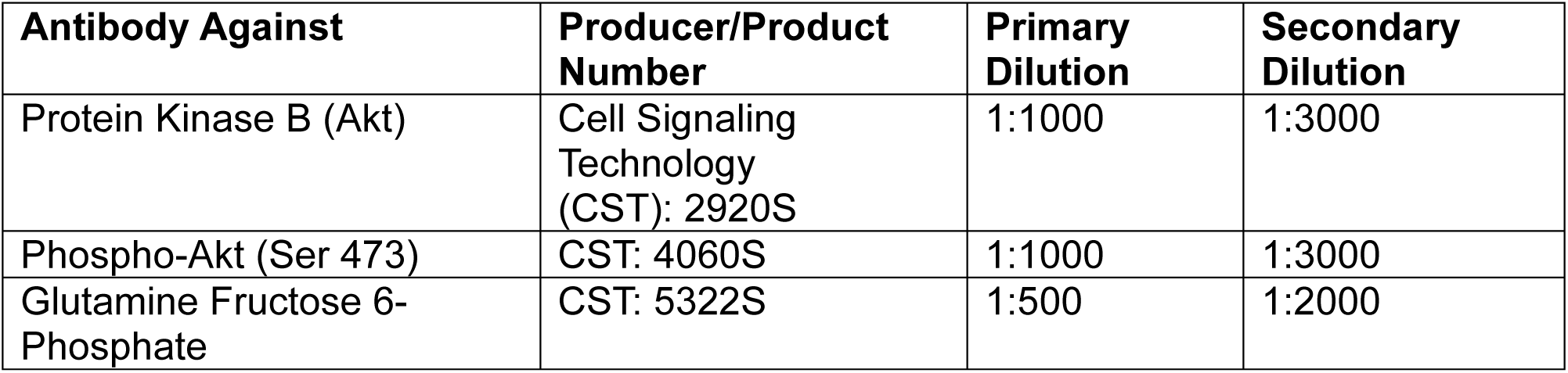
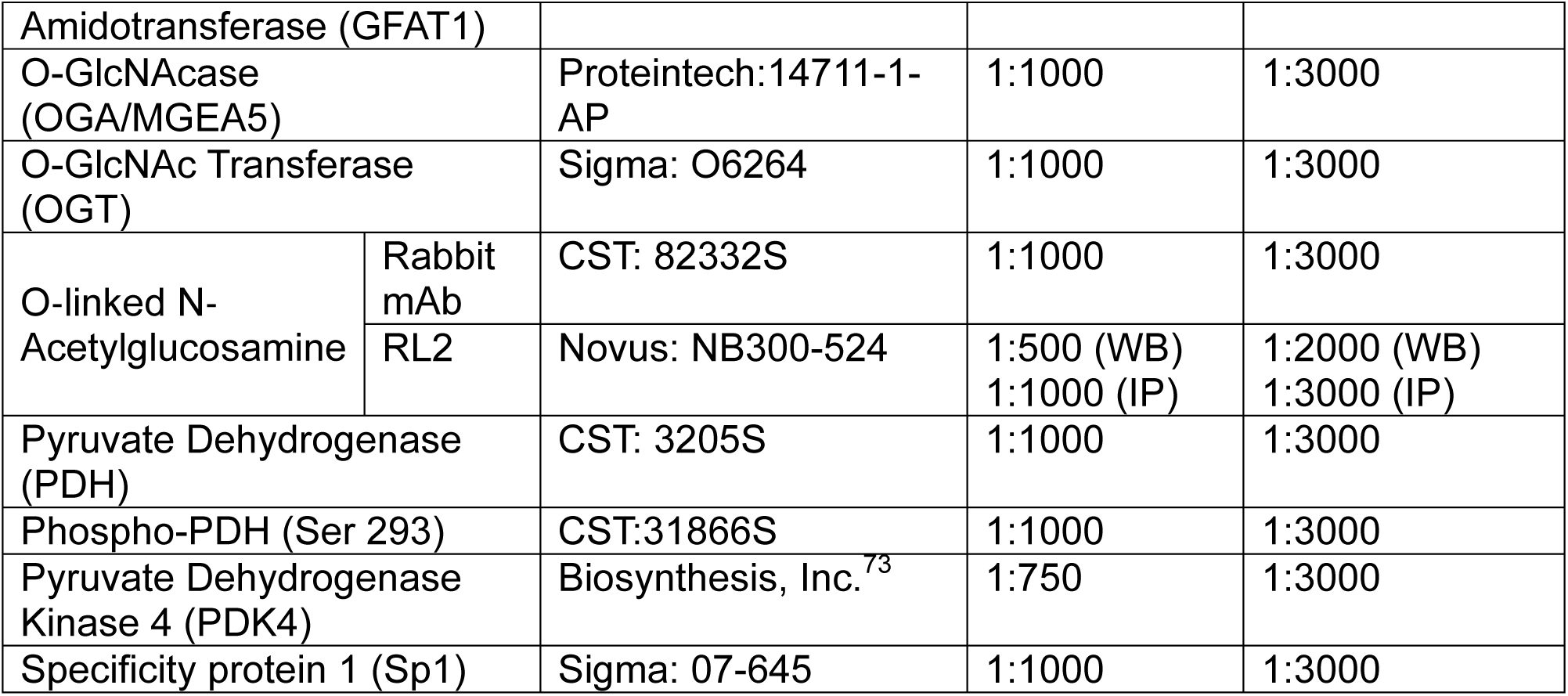
Antibodies used for Western Blot Analysis

| **Antibody Against** | **Producer/Product Number** | **Primary Dilution** | **Secondary Dilution** |
| --- | --- | --- | --- |
| Protein Kinase B (Akt) | Cell Signaling Technology (CST): 2920S | 1:1000 | 1:3000 |
| Phospho-Akt (Ser 473) | CST: 4060S | 1:1000 | 1:3000 |
| Glutamine Fructose 6-Phosphate | CST: 5322S | 1:500 | 1:2000 |

|  |  |  |  |  |
| --- | --- | --- | --- | --- |
| Amidotransferase (GFAT1) |  |  |  |  |
| O-GlcNAcase (OGA/MGEA5) |  | Proteintech:14711-1-AP | 1:1000 | 1:3000 |
| O-GlcNAc Transferase (OGT) |  | Sigma: O6264 | 1:1000 | 1:3000 |
| O-linked N-Acetylglucosamine | Rabbit mAb | CST: 82332S | 1:1000 | 1:3000 |
|  | RL2 | Novus: NB300-524 | 1:500 (WB)<br>1:1000 (IP) | 1:2000 (WB)<br>1:3000 (IP) |
| Pyruvate Dehydrogenase (PDH) |  | CST: 3205S | 1:1000 | 1:3000 |
| Phospho-PDH (Ser 293) |  | CST:31866S | 1:1000 | 1:3000 |
| Pyruvate Dehydrogenase Kinase 4 (PDK4) |  | Biosynthesis, Inc.<sup>73</sup> | 1:750 | 1:3000 |
| Specificity protein 1 (Sp1) |  | Sigma: 07-645 | 1:1000 | 1:3000 |

For immunoprecipitation assays, 400 μL TSD (50mM Tris at pH 7.5, 1% SDS, 5mM dithiothreitol) was added to the entirety of the crude nuclear pellet saved from the first (550×g) spin of total heart homogenate, and boiled at 95°C for 10 minutes, vortexing half way through. The mixture was then microcentrifuged briefly at room temperature to remove insoluble debris, and 200 μL of the supernatant was added to 1 mL ice cold TNN (50mM Tris, pH7.5, 250mM NaCl, 5mM EDTA, 0.5% NP40) with protease inhibitor. After mixing, an aliquot of this dilution was saved as a starting point control, and 4 μL Specificity Protein 1 (Sp1) antibody (Sigma: 07-645) added to 400 μL of this dilution.

After co-incubation overnight with rotation at 4°C, 60 μL washed Protein G beads were added, and incubation was repeated for 24 hours at 4°C. Following the second overnight incubation, an aliquot of the unbound supernatant was saved as a flowthrough control. Beads were washed 3 times with 1 mL ice cold TNN. Residual TNN was removed, and the beads were boiled at 95°C for 5 minutes with 20 μL TNN and 20 μL 4x LDS. Resulting immunoprecipitation samples, as well as respective starting point and flowthrough controls, were run together on a gel per the above western blot protocol.

Blots were probed with primary antibodies against Sp1 (Sigma, 07-645) and O-GlcNAc (RL2; Novus, NB300-524), followed by their respective secondary antibodies.

## Plasma Insulin Measurement

Blood was collected by cardiac puncture prior to perfusion and excision and immediately transferred to microtainer tubes on ice (BD, 365992). Blood was then centrifuged at 3000×g for 15 minutes at 4°C. The supernatant (plasma) was collected and snap frozen. Plasma insulin was measured using an ALPCO Insulin Rodent (Mouse/Rat) Chemiluminescence ELISA (80-INSMR-CH01) per the manufacturer instructions, quantified with a Cytation 5 (Agilent) in luminescence plate reading mode, and analyzed with BioTek Gen5 Microplate Reader and Imager Software (Version 3.14, Agilent). From the resulting data, as well as blood glucose measurements, Homeostatic Model Assessment for Insulin Resistance (HOMA-IR) calculations were performed as HOMA IR=[plasma insulin (pM)]×[blood glucose (mM)]/135, derived from^17,18^ HOMA-IR=[plasma insulin (μU/mL)]×[blood glucose (mM)]/22.5.

## Glycogen and TAG Liver Assays

Livers were snap frozen and powdered for use in glycogen and triglyceride (TAG) measurement assays. Liver TAG (10010303, Cayman Chemical) and glycogen (MAK465, Sigma-Aldrich) were measured using colorimetric kit-based assays according to the manufacturer instructions. Absorbance measurements at respective wavelengths were performed by Cytation5 plate reading (Agilent) and analyzed with BioTek Gen5 Microplate Reader and Imager Software (Version 3.14, Agilent).

## Statistical Analysis

For instances where only 2 groups were compared at once, data were analyzed using a two tailed, unpaired Student *t* test using GraphPad Prism (Version 10). For data sets comparing both genotype and fasting effects, a 2-way ANOVA with a main effects model was used, and Šídák’s multiple comparisons test was used for post-hoc analysis. For data sets comparing genotype, fasting, and adrenergic stress effects, a 3-way ANOVA was used with Šídák’s multiple comparisons test again used for post-hoc analysis. All data are presented as mean±SD unless otherwise indicated. Statistical significance was established at p≤0.05 and indicated in each figure legend. Principal component analysis (PCA) was performed with PCA plotter V1.02 (https://scienceinside.shinyapps.io/mvda/). Metabolomics data were analyzed with MetaboAnalyst 6.0,^19^ as described in the figure legends.

## Results

### Contribution of PFKFB2 to Mitochondrial Metabolic Flexibility

PFKFB2 regulates the rate of glycolysis, which is a key component of metabolic flexibility. We have also previously shown that altered regulation of the phosphofructokinase nexus can influence mitochondrial substrate preference^11,20^. We sought to further understand the role of PFKFB2 in downstream mitochondrial metabolic flexibility, or metabolic substrate switching, in response to 12 hours of fasting. Pyruvate dehydrogenase (PDH) is the rate limiting step of glucose oxidation and its activity is known to decrease with fasting^21,22^. We have previously demonstrated that PDH activity is increased when the phosphofructokinase regulatory nexus is disrupted by knockout of PFKFB2^11^. In mitochondria from CON hearts, PDH activity was 37.9% lower in the fasted relative to the fed group (p=0.0105; Figure 1A) and subsequently, pyruvate-supported mitochondrial state 3 respiration (the ADP-dependent maximal respiration rate; p=0.0242; Figure 1B) was 15.4% lower in fasted hearts. However, PDH activity and pyruvate-supported respiration had an attenuated response to fasting in mitochondria from cKO hearts (Figure 1A and 1B). Differences in activity were not due to abundance of total PDH, which were unchanged with fasting in CON and cKO mice (Figure 1C). There was, however, an increase in inhibitory phosphorylation of PDH in response to fasting in hearts of both genotypes (Figure 1D). Furthermore, the abundance of pyruvate dehydrogenase kinase 4 (PDK4), the predominant PDH kinase isoform in the heart, was increased in response to fasting in both genotypes (3.72-fold and 3.57-fold increases in CON and cKO mice, respectively; Figure 1E). Nonetheless, the changes in PDH activity and respiration point to a decreased capacity for glucose metabolic flexibility in mitochondria from hearts lacking PFKFB2. Though, it is interesting to note that glucose metabolism profiles of fasted cKO hearts largely recapitulate those of fed CON mice, suggesting that fasting mediates normalization of changes which occur secondary to PFKFB2 loss.

**Figure 1.**
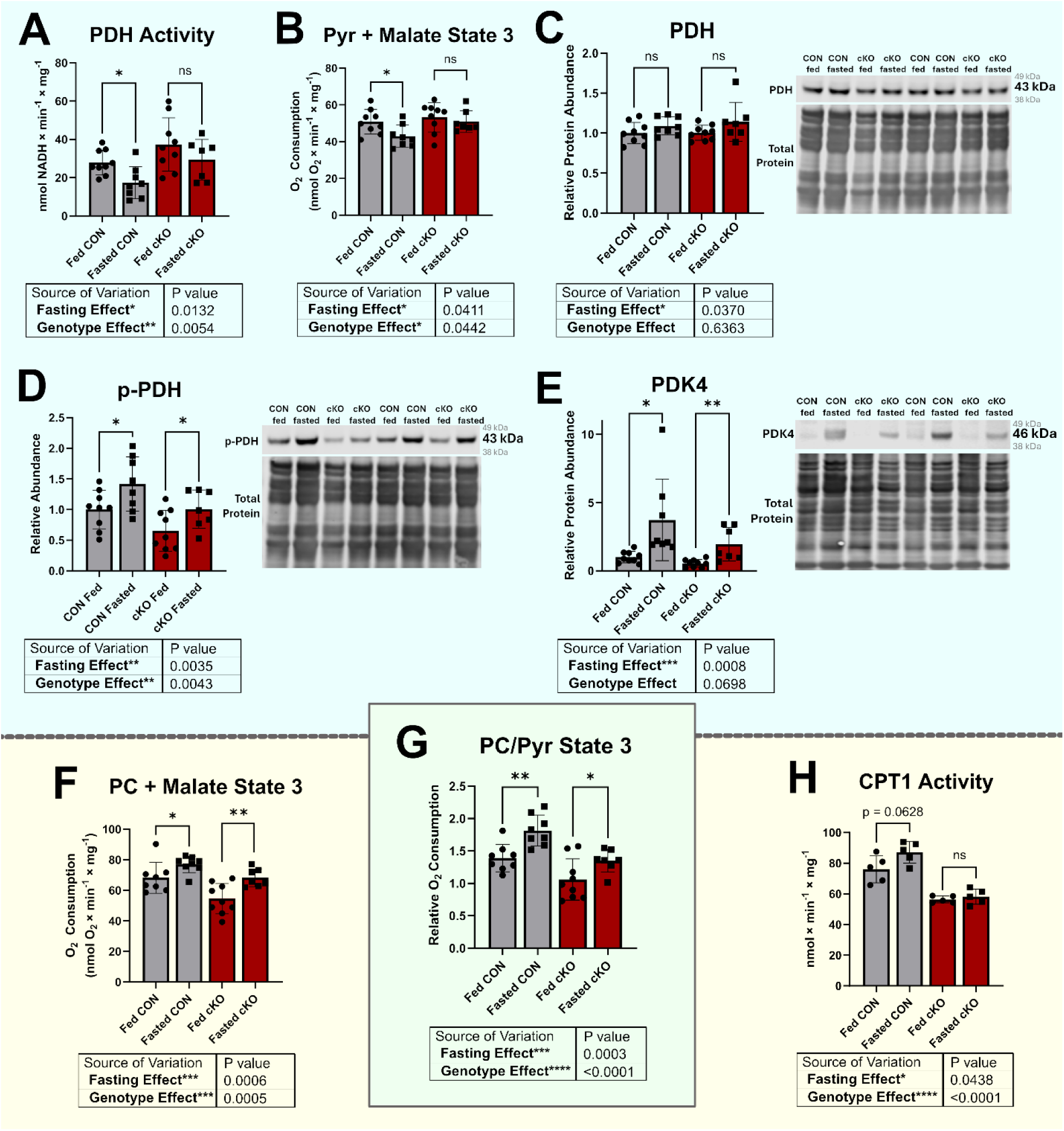
PFKFB2 loss moderately impacts cardiac mitochondrial metabolic flexibility. **A,** PDH enzyme activity was measured across groups in isolated cardiac mitochondria from fed and fasted, control (CON) and PFKFB2 knockout (cKO) hearts (n=7-9). **B,** Pyruvate (Pyr)- and malate-dependent mitochondrial respiration was measured as oxygen consumption in the ADP-dependent state (state 3; n=7-9). **C-D,** Relative abundance of total pyruvate dehydrogenase (PDH; **C**) and phosphorylated pyruvate dehydrogenase (p-PDH; **D**) were measured by western blot in the mitochondrial fraction of hearts (n=7-9). **E,** Pyruvate dehydrogenase kinase 4 (PDK4) abundance was measured by western blot in whole heart homogenate (n=7-9). **F,** Palmitoyl carnitine (PC) and malate-dependent respiration was measured in state 3 (n=7-9). **G,** The ratio of PC/Pyr-dependent State 3 respiration was calculated using measures from **B** and **F** (n=7-9). A statistical outlier in PC and malate-dependent respiration was detected in the fed CON group via a ROUT test with Q=1% and excluded in panels F and G. **H,** Carnitine palmitoyl transferase 1 (CPT1) activity was measured in isolated mitochondria(n=5). Statistics were performed as two-way ANOVA. Individual comparisons were also performed between the fed and fasted states within genotype groups by unpaired Student *t* test. Data are shown as mean ± SD. Not significant (ns) indicates *p*>0.05. \**p*≤0.05, **p≤0.01, \*\*\**p*≤0.001.

As glucose and fatty acid oxidation activities are inversely regulated, as described by the Randle cycle^23,24^, we also measured palmitoyl carnitine (PC)-supported state 3 respiration. Mitochondria from both genotypes exhibited increases in maximal PC-supported oxygen consumption when comparing fasted to fed states (p=0.0061-0.0483; Figure 1F). This is consistent with a shift toward fatty acid oxidation in the fasted state. As a metric of overall metabolic substrate preference, we took the ratio of the State 3 mitochondrial respiratory rate when supported by PC to the rate when supported by pyruvate^11,20,25,26^. We observed comparable increases in PC-relative to pyruvate-dependent respiration in CON (30.6% increase with fasting, p=0.0021) and cKO (27.5% increase with fasting, p=0.0473) mice (Figure 1G). However, because PC bypasses the rate-limiting step of fatty acid uptake, catalyzed by carnitine palmitoyl transferase 1 (CPT1), we also measured CPT1 activity directly. We observed a trend towards an increase (14.4%, p=0.0628) in CON mice with fasting but no difference in cKO mice (Figure 1H), suggesting the potential for differences in flexibility of fatty acid metabolism between genotypes that is not captured in the respirometry data.

We further investigated metabolic flexibility by performing semi-targeted metabolomics using a combination of GC-MS and LC-MS. A total of 50 metabolites were quantified (Table S1). Somewhat surprisingly, global changes in metabolism based on genotype and fed/fasted state were modest. Principal component analysis (PCA) revealed overlap between groups (Figure S1) and 2-way ANOVA with false discovery rate (FDR) revealed that 4 metabolites were significantly affected by either diet or genotype (Figure 2A and S1). The ketone body, 3-hydroxybutyrate, was significantly affected by fasting, as expected. However, upon correction for multiple comparisons, this only reached statistical significance in CON mice. By genotype, aspartate and serine, two glucose-derived amino acids, were significantly elevated in cKO relative to CON mice (Figure 2A). This increase was driven largely by cKO hearts in the fed state, with fasting considerably mitigating the increase in cKO mice (Figure 2A). Succinic acid was affected by both genotype and fasting, though no differences were observed in succinate-dependent mitochondrial respiration across groups (Figure S2). Despite the modest statistically significant differences among the 50 metabolites, there were several strong correlations with both genotype and fasting. Figure 2B shows metabolites that were positively or negatively correlated with cKO genotype (*left*) and fasting (*right*).

**Figure 2.**
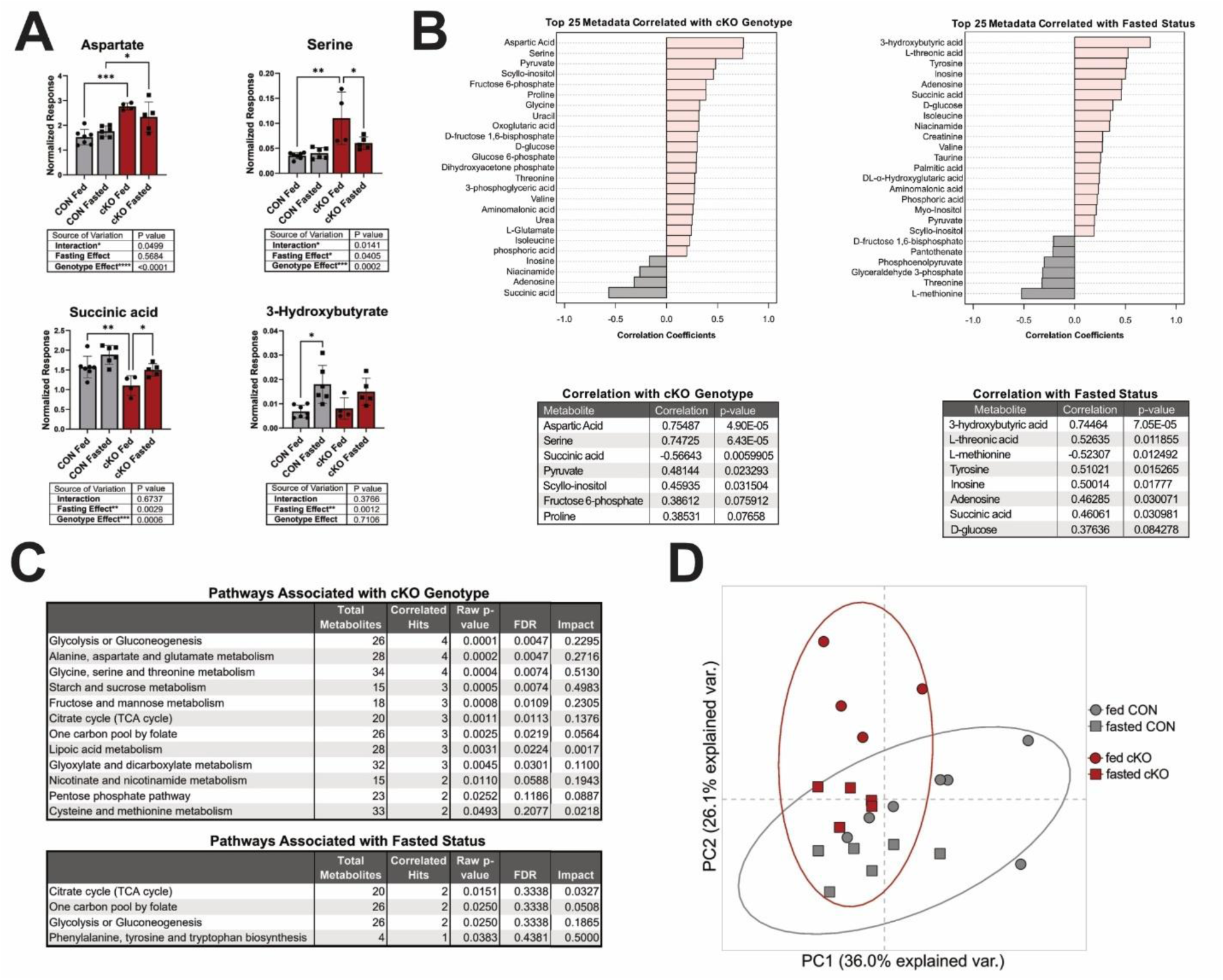
Key Metabolite Changes and Correlations Associated with PFKFB2 Loss and Fasting. Metabolites were measured using a combination of liquid chromatography-mass spectrometry (LC-MS) and gas chromatography-mass spectrometry (GC-MS) on PFKFB2 knockout (cKO) or control (CON) hearts (n=4-7). **A,** Metabolites which were significantly affected by fasting include Aspartate (LC-MS), Serine (GC-MS), Succinic Acid (GC-MS), and 3-Hydroxybutyrate (GC-MS). Data are shown as mean ± SD. **B,** Metabolites which are significantly correlated with cKO genotype (left, n=9-13) or fasted status (right, n=11) are shown, with those metabolites whose correlation achieves statistical significance of p<0.10 listed in tables below. **C,** Pathways correlated with cKO genotype or fasted status with statistical significance of p<0.05 are listed. **D,** Principal component analysis on metabolites with correlation significance of p<0.10 (n=4-7). Circles indicate fed status, squares indicate fasted status, grey indicates control, and red indicates cKO. A, Metabolites were analyzed by two-way ANOVA with Sidak’s post hoc test for multiple comparisons between groups which varied by a single factor. \**p*≤0.05, **p≤0.01, ***p≤0.001, ***p≤0.001. B-C, Correlations and pathway analyses were performed with MetaboAnalyst 6.0 using default settings.

Pathway analysis on correlated metabolites revealed genotype-dependent upregulations in fructose metabolism as well as glycolysis and gluconeogenesis in cKO mice (Figure 2C), consistent with our previous report^11^. Amino acid related pathways, including alanine, aspartate, and glutamate metabolism, in addition to phenylalanine, tyrosine, and tryptophan biosynthesis, were upregulated both in response to cKO genotype and fasted status (Figure 2C). Lastly, we performed PCA on significantly (p<0.1) correlated metabolites which revealed clear separation by group (Figure 2D).

Interestingly, the fed cKO hearts grouped distinctly from other groups. However, fasted cKO hearts grouped much more similarly with CON mice of both groups. This supports that many of the metabolic changes driven by PFKFB2 loss are ameliorated in the fasted state.

## Coordination Between the PFKFB2/PFK-1 Nexus and Substrate Availability Impacts O-GlcNAc Regulation

We have previously shown that the loss of PFKFB2 leads to an increase in the post-translational modification O-GlcNAcylation^11^. A plausible driver of this increase is that disruption of the phosphofructokinase nexus promotes fructose 6-phosphate spillover into the hexosamine biosynthesis pathway (HBP), which provides substrate for O-GlcNAcylation. Therefore, we hypothesized that during fasting, the decrease in glucose metabolism may mitigate this increase. Consistent with our previous report, we observed a 42% increase in O-GlcNAcylation in hearts of fed cKO relative to fed CON mice (p=0.0009; Figure 3, S3A). This increase in O-GlcNAcylation in cKO hearts was partially ameliorated by fasting (Figure 3, S3A). No impact on O-GlcNAcylation levels was observed in CON mice with fasting. Because most antibody-detected O-GlcNAc substrates are approximately 50 kDa or larger, we next separately quantified O-GlcNAcylation as only modified proteins above 50 kDa. When quantified via this approach^27^, a 66% increase in O-GlcNAcylation levels was observed in fed cKO relative to fed CON mice (p=0.0004; Figure 3, S3B) and O-GlcNAcylation levels were significantly decreased in fasted cKO mice relative to fed (p=0.0280; Figure 3, S3B).

**Figure 3.**
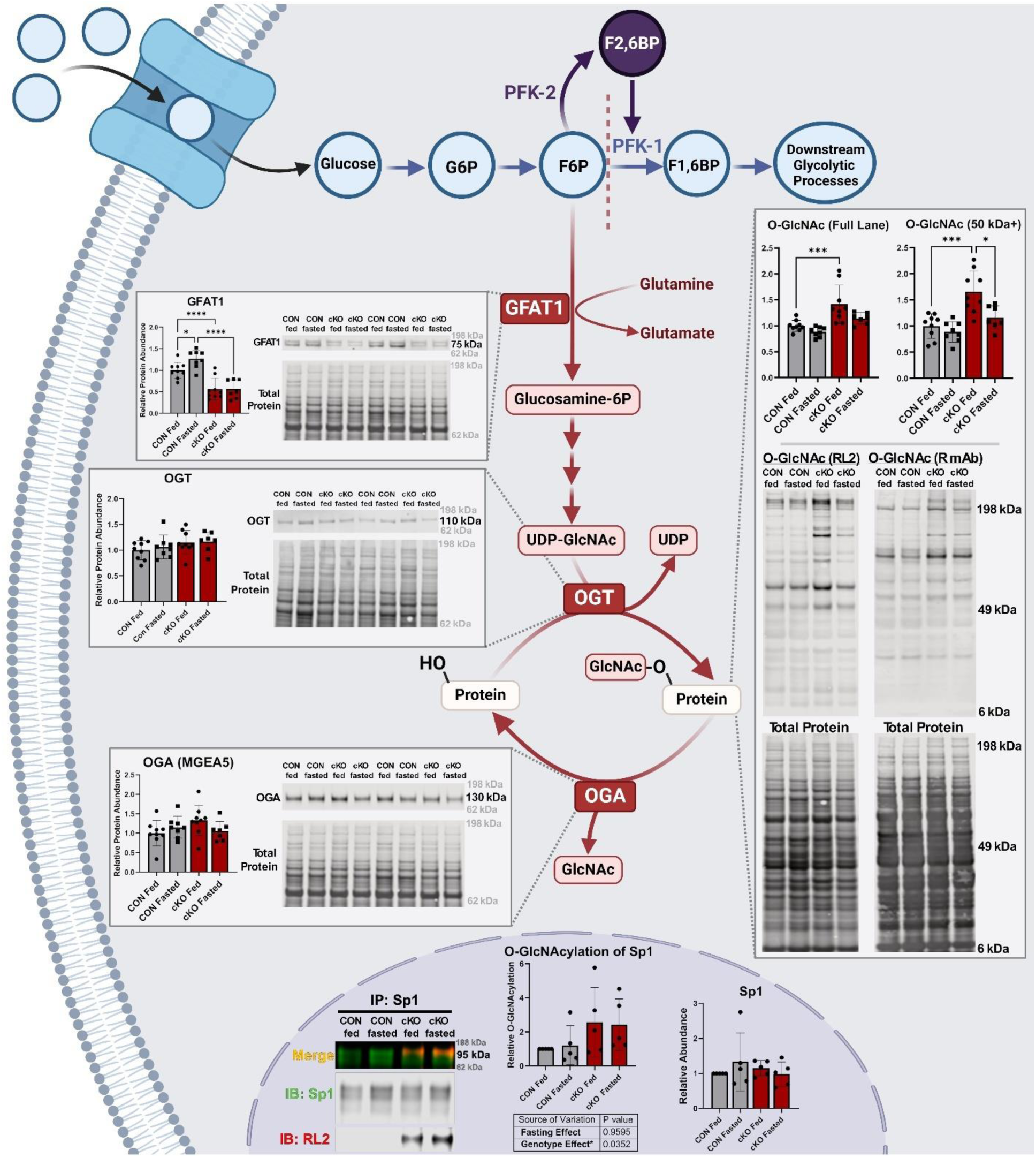
O-GlcNAcylation is increased in fed, but not fasted, cKO hearts, suggesting substrate-level regulation. Glutamine-fructose 6-phosphate amidotransferase (GFAT1), O-GlcNAc transferase (OGT), O-GlcNAcase (OGA (MGEA5)), and overall protein O-GlcNAcylation were measured by western blot, using heart homogenate from fed and fasted, control (CON) and PFKFB2 knockout (cKO) hearts (n=7-9). O-GlcNAc levels on western blot were measured using the average of signals from two different pan-specific antibodies targeting the modification (mouse monoclonal (RL2 clone) and rabbit monoclonal (R mAb)), and quantified either as the full lane of proteins (left) or only proteins which are 50 kDa or larger (right). O-GlcNAcylation of specificity protein 1 (Sp1) was measured by immunoprecipitation of Sp1, subsequently blotting for both Sp1 and O-GlcNAc (RL2 antibody), and taking the quotient. Sp1 was quantified directly, and the ratio of RL2 signal/Sp1 signal was used to determine the relative O-GlcNAcylation of Sp1 (n=5). Data were analyzed by two-way ANOVA with Sidak’s post-hoc test for multiple comparisons between groups which differed by one factor. \**p*≤0.05, ***p≤0.001, ****p≤0.0001. Data are shown as mean ± SD.

Interestingly, this appears to be driven largely by cytosolic modification as O-GlcNAcylation is unchanged across groups in the mitochondrial fraction, except for a slight increase in O-GlcNAcylation levels in fasted cKO relative to CON mitochondria (Figure S4).

While a possible explanation for the attenuation of O-GlcNAcylation is decreased glucose uptake and availability in the heart in the fasted state, O-GlcNAcylation can also be affected by the abundance and regulation of HBP enzymes. The rate limiting step of the HBP is catalyzed by glutamine-fructose 6-phosphate amidotransferase (GFAT). We observed a 26% increase in GFAT1 abundance with fasting in CON mice (p=0.0138). Unexpectedly, we observed a significant decrease (44%) in GFAT1 abundance in cKO mice relative to fed CON (p<0.0001; Figure 3) regardless of fed/fasted status.

Transcription of *Gfpt1*, the gene encoding GFAT1, is regulated in part by the transcription factor specificity protein 1 (Sp1)^28^. Further, Sp1 can be inactivated by O-GlcNAcylation in a glucose availability-dependent manner^29–31^. We hypothesized that the decrease in GFAT1 levels could be attributable to a feedback response via increased Sp1 O-GlcNAcylation. We therefore measured the abundance and relative O-GlcNAcylation of Sp1 by immunoprecipitation in nuclear-enriched fractions (Figure 3, S5). While Sp1 levels were unchanged based on fed or fasted state, there was a genotype-dependent increase (p=0.0352) in O-GlcNAcylation of Sp1 in cKO hearts (Figure 3). This points to a potential for compensatory suppression of Sp1-mediated transcription of *Gfpt1* which could explain the decrease in GFAT1 protein abundance in cKO mice.

O-GlcNAcylation is also regulated at the levels of its addition and removal by O-GlcNAc Transferase (OGT) and O-GlcNAcase (OGA, or MGEA5) respectively. However, abundance of both OGA and OGT was unchanged across groups (Figure 3).

Collectively, these data point to a role of glucose availability in driving increased O-GlcNAcylation in the context of PFKFB2 loss. This supports that a dynamic coordination between glucose availability and the phosphofructokinase nexus may regulate levels of protein O-GlcNAcylation.

## The Role of PFKFB2 in Cardiac Insulin Response and Whole-Body Glucose Homeostasis

We next sought to determine if the loss of cardiac PFKFB2 affected cardiac insulin signaling and whole-body glucose homeostasis. Consistent with our previous report^11^, the abundance of Akt, the primary intracellular signal transducer for insulin signaling, and it’s activated form, p-Akt (Ser 473) were increased in fed cKO relative to fed CON hearts (p=0.022-0.023; Figures 4A, 4B, and 4D). The increase in p-Akt in cKO relative to CON hearts also occurred under fasted conditions (p=0.0027; Figure 4B and 4D). As expected with fasting, p-Akt decreased 32% in CON mice (p=0.0084; Figure 4B and 4D). In cKO hearts, there were fasting-induced decreases in both Akt and p-Akt, though these failed to reach statistical significance. Furthermore, fasting returned Akt and p-Akt levels in cKO hearts to approximately those of fed CON hearts, consistent with the overall trend of glucose utilization changes in the cKO heart being normalized by fasting. Due to comparable trends in abundance of total Akt and p-Akt, differences in the ratios of the two failed to reach statistical significance (Figure 4C and 4D).

**Figure 4.**
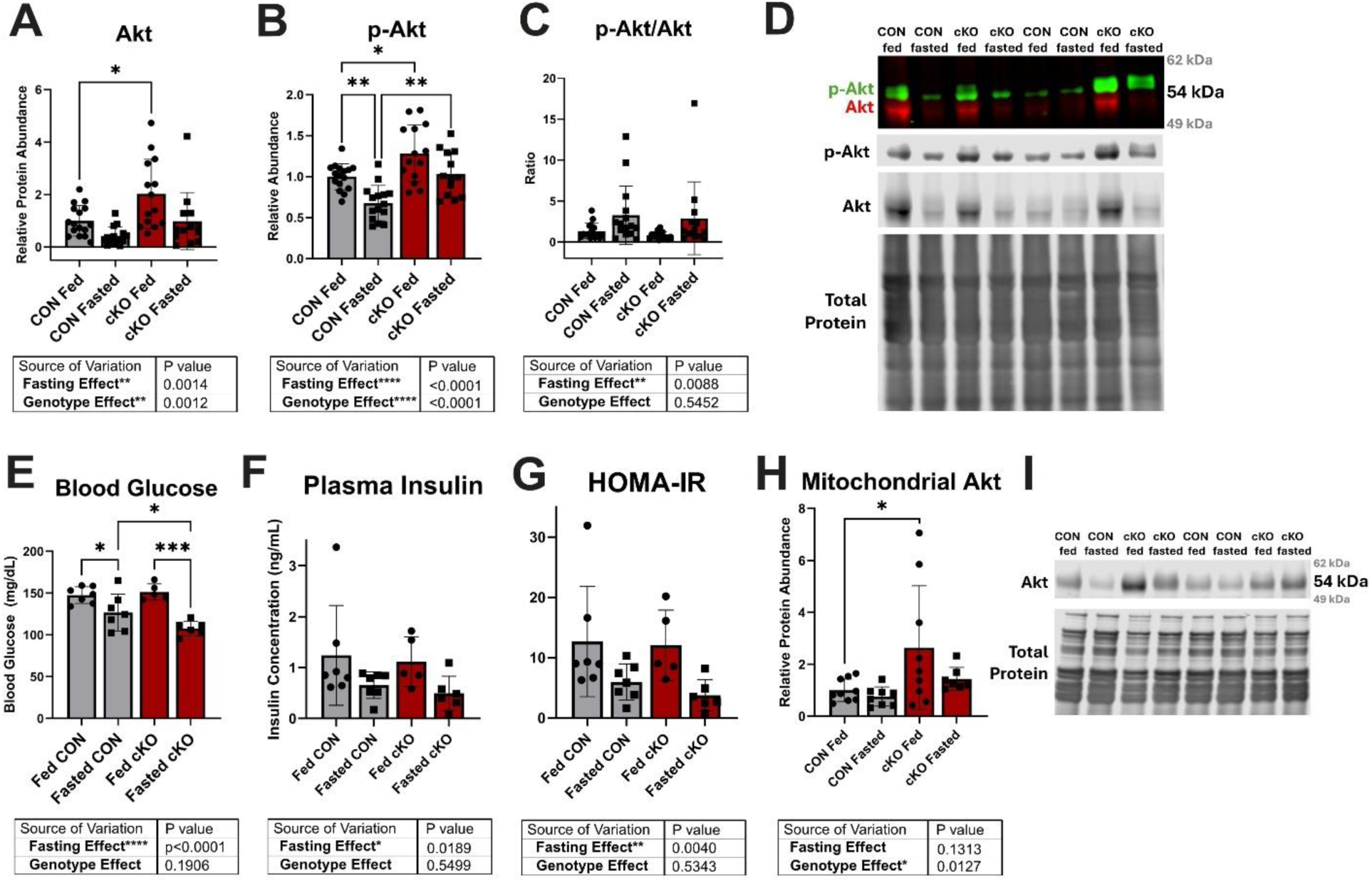
Loss of cardiac PFKFB2 enhances insulin signaling and glucose utilization in the fasted state. **A-D,** Relative abundance of total Akt (**A**), phosphorylated Akt (p-Akt, **B**), and the ratio of p-Akt/Akt (**C**) were measured by western blot in fed and fasted PFKFB2 knockout (cKO) and control (CON) hearts (n=13-16). A sample blot is shown with p-Akt in green and total Akt in red (**D**). **E,** Blood glucose was measured immediately prior to sacrifice (n=5-7). **F**, Plasma insulin was measured from blood collected by cardiac puncture. **G,** Homeostatic Model Assessment for Insulin Resistance (HOMA-IR) was calculated from blood glucose and plasma insulin measures (n=5-7). **H-I,** Total Akt was measured by western blot in mitochondrial fraction (n=7-9). Data are shown as mean ± SD. All data were analyzed by two-way ANOVA with Sidak’s post-hoc test for multiple comparisons between groups varying by one factor. \**p*≤0.05, **p≤0.01, ***p≤0.001.

At the whole-body level, we observed a 14.1% decrease in blood glucose in the fasted state in CON mice (p=0.0293). Interestingly, cKO mice showed a more significant reduction in blood glucose with fasting (28.80% decrease, p=0.0002; Figure 4E). While a decrease in circulating (plasma) insulin with fasting failed to reach statistical significance in mice of either genotype, there were trends toward decreases, and a statistically significant main fasting effect was observed by 2-way ANOVA (Figure 4F). A commonly used metric of insulin sensitivity is the Homeostatic Model Assessment for Insulin Resistance (HOMA-IR). A lower value is a proxy for increased insulin sensitivity.

We observe trends toward decreases in HOMA-IR in both CON (p=0.2431) and cKO (p=0.2193) mice with fasting (Figure 4G) with minimal difference by genotype. The main effect of fasting reached statistical significance (p=0.0040). HOMA-IR is a useful measure of whole-body glucose homeostasis. However, in the fasted state with abated glucose consumption and insulin secretion, a lower HOMA-IR may not accurately indicate higher insulin sensitivity than in the fed state and should be interpreted cautiously.

Lastly, it has been previously demonstrated that Akt associates with the mitochondria and subsequently activates PDH^11,32^. Therefore, given our changes in pyruvate metabolism in the fasted state, we measured Akt abundance in the mitochondrial fraction. We found a significant genotype main effect, driven mostly by a 2.64-fold increase in mitochondria-associated Akt in fed cKO relative to CON mice (p=0.033). This increase was ameliorated with fasting (Figure 4H and 4I). These data further support a normalizing effect of fasting on glucose metabolism in cKO hearts.

Cumulatively, these data point to a directionality of the relationship between PFKFB2 and insulin signaling, whereby PFKFB2 loss does not provide negative feedback, but perhaps even compensatory upregulation of Akt signaling in an isolated system.

## The Role of Cardiac PFKFB2 in Systemic Response to Stress

It is established that adrenergic stress drives glucose uptake and utilization in the healthy heart. This is mediated by multiple mechanisms, including the phosphorylation of PFKFB2 by PKA^8,33^. Having shown that PFKFB2 is affecting insulin sensitivity and glucose use at baseline, we next sought to determine the impact of PFKFB2 loss on response to stress in both cardiac and systemic metabolism. To do so, we challenged mice acutely with caffeine and epinephrine, which are commonly used to test cardiac physiological stability. Mice were injected after 12 hours of fasting, and the stress was carried out for 20 minutes prior to tissue collection. Blood glucose levels were measured before fasting, as well as after fasting both pre- and post-pharmacologic stimulant stress (Figure 5A). To establish efficacy of the stress, we measured p-PKA substrate levels as PKA is indirectly activated by both epinephrine and caffeine through their independent effects on cAMP (Figure S6). Two-way ANOVA revealed main effects of both cKO genotype and stimulant stress on phosphorylated PKA substrate levels (Figure S6). The effects of stress were relatively uniform across groups, with stress leading to a 23-40% increase in phosphorylated PKA substrate levels in stressed groups relative to their respective baseline (unstressed) group (Figure S6).

**Figure 5.**
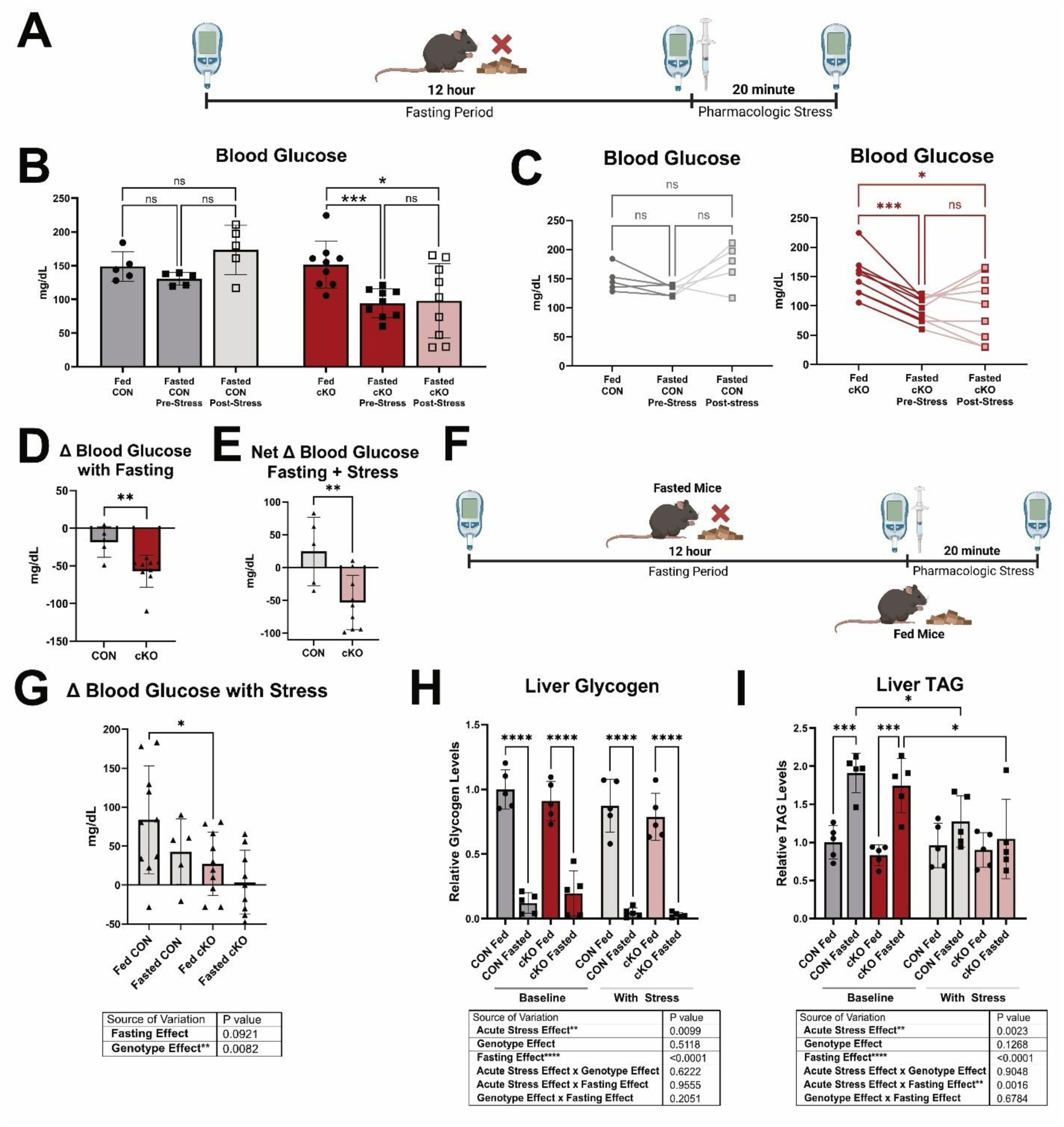
Acute stimulant stress promotes differential systemic glucose homeostasis in cKO mice. **A,** In the first group of mice, blood glucose was measured in the fed state. Mice were then fasted for 12 hours, after which blood glucose was measured a second time (fasted, pre-stress). Following the second blood glucose measurement, a stimulant stress of caffeine and epinephrine was injected intraperitoneally. 20 minutes after injection, a third blood glucose measurement was acquired (fasted, post-stress). **B-C,** Average blood glucose levels across fed, fasted pre-stress, and fasted post-stress states (**B**) are plotted alongside individual trends (**C**) in fed and fasted CON and cKO (n=5-9). **D-E,** Change in blood glucose was calculated, first with fasting alone (**D**) and then with both fasting and stimulant stress (**E**, n=5-9). **F,** In another group of mice, blood glucose was measured in the fed state. The stimulant stress of caffeine and epinephrine was then administered to fed mice and blood glucose was measured again 20 minutes post-injection. **G,** Change in blood glucose with stimulant stress in fed or fasted, CON or cKO mice is shown (n=5-10). **H-I,** Relative liver glycogen (**H**) and liver triglycerides (TAG; **I**) were measured in fed and fasted CON and cKO mice, with and without stimulant stress (n=5). Where applicable, data are displayed as mean ± SD. **B-C** were analyzed by repeated measures one-way ANOVA with Sidak’s multiple comparisons test. **D-E** were analyzed by unpaired Student’s *t* test. **G** was analyzed by two-way ANOVA with Sidak’s post-hoc test for multiple comparisons. **H-I** were analyzed by three-way ANOVA with Sidak’s post-hoc test for multiple comparisons. \**p*≤0.05, **p≤0.01, ***p≤0.001, ****p≤0.0001.

Under normal fed conditions, adrenergic stimulation leads to a mobilization of liver glycogen, increasing circulating glucose concentrations^34^. Here, we observed that in CON mice, there was a slight trend towards a decrease in blood glucose levels following 12 hours of fasting that reversed to trend toward an increase with acute stress (Figure 5B and 5C). However, cKO mice demonstrated a 37.65% decrease in blood glucose with fasting (p=0.0001) which was sustained with stimulant stress (Figure 5B and 5C). This culminated in a significantly greater decrease in blood glucose both with fasting alone (p=0.0061; Figure 5D) and with the combination of fasting and stimulant stress (p=0.0095; Figure 5E) in cKO compared to CON mice. We next tested whether this stress-induced change in blood glucose was specific to the fasted state by pharmacologically stressing fed mice at the same dose and duration, measuring blood glucose pre-and post-stress (Figure 5F). We observed that the pharmacologic stimulant stress increased blood glucose considerably in fed CON mice (84.0 mg/dL; Figure 5G) but to a lesser extent in fed cKO mice (27.3 mg/dL; Figure 5G). Similarly, there was minimal response of blood glucose levels to stress in fasted cKO mice (Figure 5G). Liver weights were significantly decreased with fasting in mice of both genotypes (Figure S7). We thus hypothesized that the tempered blood glucose response in cKO mice was due to a depletion of liver glycogen, in conjunction with increased cardiac glucose use, albeit non-canonical use through auxiliary glucose metabolism pathways in cKO hearts. While differences in liver glycogen between genotypes with stress did not reach statistical significance, there was a negligible amount of glycogen remaining in livers of stressed and fasted cKO mice (2.88% of that which is present in unstressed, fed CON mice; Figure 5H). However, glycogen levels were also depleted in livers from unstressed CON mice. This points towards a likely contribution of the rate of glucose consumption by the stressed heart in cKO relative to CON mice on circulating glucose levels. Given the changes in cardiac metabolic substrate utilization during fasting, and the heart’s predominant fatty acid dependence, we also investigated differences in liver triglycerides (TAG). Liver TAG were increased, as expected, with fasting in the unstressed state. Stimulant stress prevented this increase in liver TAG (Figure 5I). However, there were no observed differences based on genotype (Figure 5I).

Finally, given the changes in glucose levels and an established role of O-GlcNAcylation in the heart’s response to stress, we measured protein O-GlcNAc levels in hearts from fed and fasted CON and cKO mice in the presence and absence of the stimulant stress. Interestingly, we found that all groups showed an increase in O-GlcNAcylation (0.51-2.12-fold, p<0.0001-p=0.0224; Figure 6, S3C) relative to their non-acutely stressed counterparts. While O-GlcNAc levels are increased in stressed cKO mice relative to stressed CON, this increase was not ameliorated with fasting in the way that it is in the unstressed state (Figure 6, S3C). This is likely due to an increase in cardiac glucose uptake with stress, independent of post-prandial state. Together, these data highlight the heart’s ability to respond to stress, the importance of regulated glucose use in the heart, and the systemic impacts which can arise if cardiac glucose use is improperly regulated.

**Figure 6.**
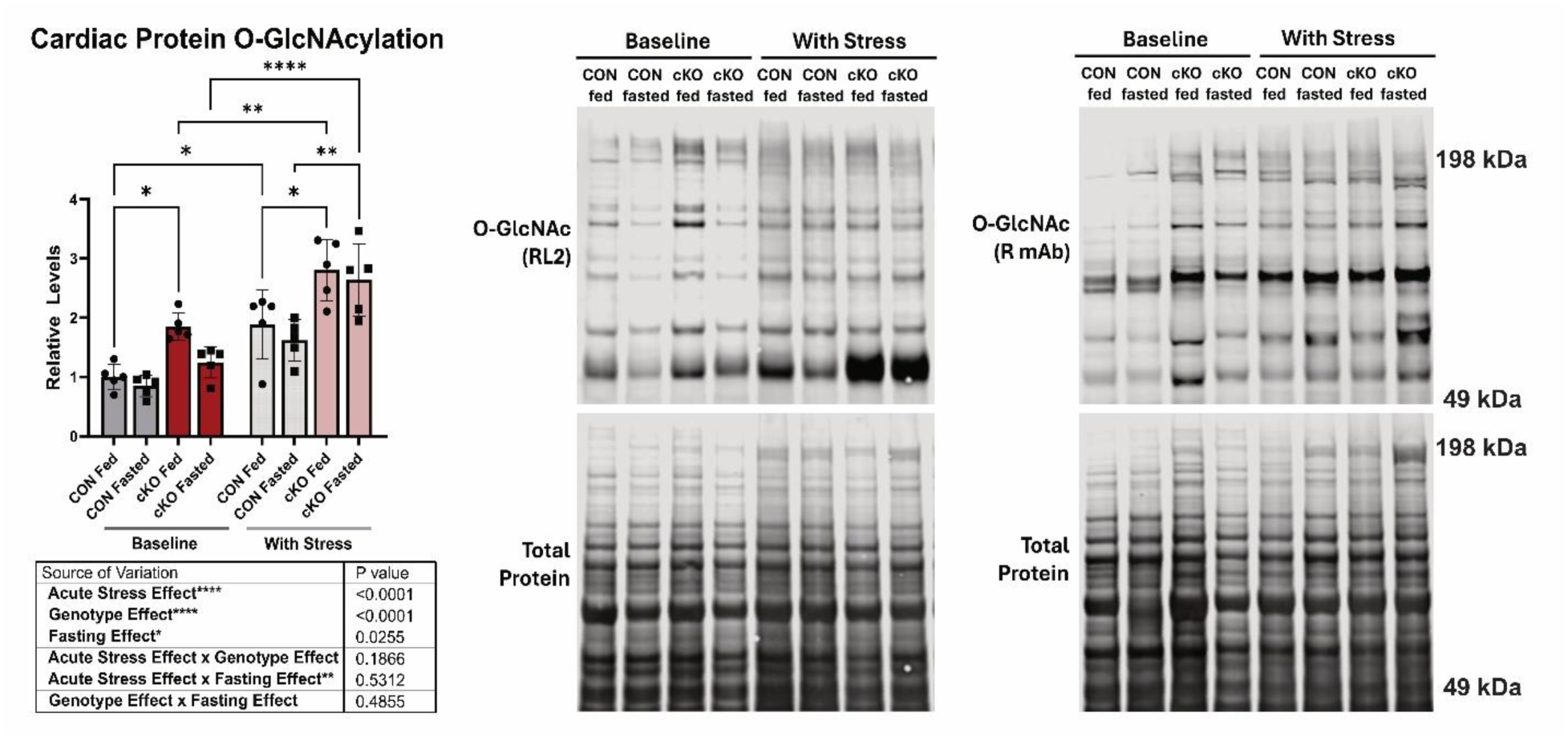
Stimulant stress drives increased cardiac O-GlcNAcylation in CON and cKO mice. Relative levels of protein O-GlcNAcylation of proteins 50 kDa or larger were measured using a pan-specific antibody in stressed (caffeine and epinephrine) and baseline (unstressed), fed and fasted PFKFB2 knockout (cKO) and control (CON) hearts (n=5). Data are shown as mean ± SD and were analyzed by three-way ANOVA with Sidak’s post-hoc test for multiple comparisons. \**p*≤0.05, **p≤0.01, ****p≤0.0001.

## Discussion

The role of PFKFB2 in metabolic flexibility is both dynamic and powerful in that its bifunctionality allows PFKFB2 to both produce and degrade the strongest allosteric activator of glycolysis, fructose 2,6-bisphosphate^6,7^. When phosphorylated in response to stress, adrenergic signaling, or insulin signaling, PFKFB2 acts as a kinase, producing fructose 2,6-bisphosphate and thereby activating glycolysis. When dephosphorylated, PFKFB2 is a phosphatase, degrading fructose 2,6-bisphosphate. This is pivotal to the heart’s ability to compensatorily upregulate glucose use in response to stress or simply glucose availability and reciprocally, to conserve glucose in the fasted state. We have previously shown that dephosphorylated PFKFB2 is degraded with decreased insulin signaling^3^. One might posit that this degradation in healthy conditions serves to maintain a baseline fructose 2,6-bisphosphate pool, as sustained levels of dephosphorylated PFKFB2 would otherwise degrade all remaining fructose 2,6-bisphosphate. While this physiological mechanism is important in short term settings such as overnight fasting, it has potential to become pathological with chronic loss of insulin signaling, such as that seen in type 1 diabetes and type 2 diabetes, which are marked by impaired insulin production and sensitivity, respectively. This pathologic potential is evidenced by our previous work, which showed that constitutive cardiac PFKFB2 loss drives structural, mechanical, and electrophysiological remodeling in the heart, as well as a sudden premature mortality^11^. This is likely attributable, at least in part, to activation of ancillary pathways including but not limited to the hexosamine biosynthesis pathway^11^.

Furthermore, we have also shown that overexpression of PFKFB2 confers protection against cardiac metabolic and functional changes in a high fat diet-induced model of metabolic syndrome^26^. Therefore, understanding the contributions of PFKFB2 loss to glucose homeostasis on more prolonged time scales becomes critical to understanding metabolic flexibility in the diabetic heart. This is particularly highlighted by contexts such as fasting where the healthy heart would normally downregulate PFKFB2 to confer flexibility and protect glucose homeostasis.

Dysregulation of, or fluctuations in, blood glucose levels with overnight fasting are not uncommon in diabetes and can pose additional challenges to glycemic control in individuals afflicted with this disease. This is highlighted by greater incidence and risk of nocturnal hypoglycemia in individuals on insulin therapy^35^. Reciprocally, the dawn phenomenon can lead to hyperglycemic spikes when an early morning rise in glucose is not met with a sufficient increase in peripheral insulin signaling^36–38^. This can drive increased intraday circulating glucose variability^37^. These factors, among others, contribute to the many cardiovascular risks associated with nocturnal fasting which are specific to, or accentuated in, individuals with diabetes. An example of these impacts is non-dipping, the failure of systolic blood pressure to decrease overnight or worse, reverse dipping, a tendency for blood pressure to increase overnight^39,40^. Further, arrhythmia and syndromes such as dead in bed syndrome, an unexplained overnight mortality, are linked to glucose control and specifically nocturnal hypoglycemia^41–43^.

Conversely, it has been suggested that intermittent fasting may confer benefit in individuals with diabetes as well as healthy individuals^10,44,45^. This dichotomy underscores the need to better understand how the body’s response to fasting is affected by the ongoing loss of PFKFB2 in the hearts of individuals with diabetes.

A hallmark of pathology in the diabetic heart is metabolic inflexibility. While PFKFB2 is important in up- or down-regulation of glycolysis, how it contributes to downstream mitochondrial metabolic flexibility in response to nutrient availability is unknown. Here, we found that there was a moderate impairment in flexibility of mitochondrial glucose oxidation and pyruvate-dependent respiration (Figure 1A-B). With regard to cardiac mitochondrial fatty acid use, as measured by palmitoylcarnitine (PC)-dependent respiration, flexibility with fasting appeared to be preserved (Figure 1F, 1H). Though, a potential caveat to this approach is that by providing PC directly to mitochondria, we bypass the key regulatory step of acyl-carnitine production on the outer mitochondrial membrane at the point of carnitine palmitoyl transferase I (CPT1), which is necessary for long chain fatty acid import into the mitochondria. When we directly measured CPT1 activity we observed no changes in activity between the fed and fasted states in cKO mice. This suggests that the respirometry data may overestimate the metabolic flexibility of cKO hearts and that fatty acid metabolic flexibility of the cKO hearts may also be impaired.

Interestingly, when evaluating values of mitochondrial metabolic parameters, most genotype-dependent changes observed in the fed cKO were normalized with fasting, thereby mimicking fed CON mice (Figure 1B-D, 1F-G). For example, glucose oxidation, which is likely supported in the cKO heart by shunting of upstream glycolytic intermediate into the pentose phosphate pathway and then back into glycolysis downstream of the phosphofructokinase nexus^11^, increased in fed cKO mice. However, glucose oxidation levels in fasted cKO mice were comparable to those observed in fed CON mice. Many whole-cell level changes related to glucose use which were observed in response to PFKFB2 loss in the fed state were also normalized by fasting.

Abundance of the glucose-derived amino acids serine and aspartate increased in the fed cKO, but this was ameliorated with fasting (Figure 2A). Of note, the opposite effect is observed in a kinase-dominant cardiac PFK-2 overexpression model, where aspartate levels increased with fasting^46^. This points to a role of phosphofructokinase nexus disruption in affecting glucose metabolism-associated processes, specifically when glucose availability is sufficient. In support of this principle, principal component analysis showed that fed cKO hearts grouped distinctly from other groups (Figure 2D), with fasted cKO hearts grouping more similarly to CON mice of both states.

Another aspect of glucose metabolism that was increased in the fed cKO heart but normalized by fasting was the total level of the post-translational modification O-GlcNAcylation (Figure 3). Work from the Chatham and Jones labs has shown that O-GlcNAc can serve cardioprotective roles under conditions of short-term stress^47–53^.

However, sustained increases in protein O-GlcNAcylation have pathological implications in the heart including promotion of cardiac dilation, functional and metabolic impairments, and increased propensity for premature mortality^14,27,54^. Further demonstrating the relevance of its chronic upregulation to cardiac pathology, this post-translational modification is upregulated in both heart failure and diabetic heart disease^14,54–57^.

The regulation of O-GlcNAcylation is complex. O-GlcNAcylation occurs when an N-acetylglucosamine group from uridine diphosphate N-acetylglucosamine (UDP-GlcNAc) is transferred to serine, threonine, and tyrosine residues of target proteins^58^. The supply of this UDP-GlcNAc arises from the hexosamine biosynthesis pathway, which integrates nutrient sensing from glucose, amino acid metabolism (glutamine), fatty acid metabolism (acetyl CoA), and nucleotide metabolism (uridine triphosphate). Increased glucose uptake and utilization through GLUT4 overexpression enhances mitochondrial protein O-GlcNAcylation in hyperglycemia, supporting glucotoxicity and impaired mitochondrial function^30^. However, seemingly contrasting work points to a striking increase in O-GlcNAc levels with glucose deprivation in in vitro studies, highlighting the gaps in our understanding of the substrate-level regulation of this post-translational modification^59^. Here, we tested a hypothesis that O-GlcNAcylation levels can be regulated by coordination between phosphofructokinase activity and glucose substrate availability.

The Hill laboratory has previously performed ^13^C_6_-glucose-based glycolytic flux assays in primary neonatal rat ventricular myocytes, using models which manipulate PFK-1 activity through kinase-dominant or phosphatase-dominant forms of PFK-2 and thus alteration of fructose 2,6-bisphosphate levels^60^. Through these tracer studies, they have shown that the phosphofructokinase nexus serves as an important regulatory point for the hexosamine biosynthesis pathway (HBP). Decreasing fructose 2,6-bisophosphate levels with a phosphatase-dominant PFK-2 overexpression promoted flux through ancillary pathways. Reciprocally, increasing PFK-2 kinase activity decreased HBP flux^60^. This suggests that when glycolytic flux through the phosphofructokinase nexus is impeded, a spillover occurs into the hexosamine biosynthesis pathway. Consistent with cardiac PFKFB2 being kinase-predominant ^61^, PFKFB2 knockout hearts have an increase in O-GlcNAc levels in the fed state, which more closely aligns with the metabolic phenotype of the phosphatase-dominant PFK-2 overexpression model.

Finally, PFKFB2 cKO mice also demonstrate cardiac dilation, cardiac functional impairment, and sudden premature death^11^ which are also observed in models of increased O-GlcNAc^14,27^.

Unlike other post-translational modifications, such as phosphorylation which is regulated by numerous kinases and several phosphatases, O-GlcNAc moieties are added by a single “writer”, O-GlcNAc transferase, and “eraser”, O-GlcNAcase. We therefore measured abundance of these enzymes and found no difference by fed/fasted status or genotype (Figure 3). While measuring the abundance of O-GlcNAc regulatory enzymes is not a comprehensive means of evaluating regulation of the respective steps, it suggests a reduced possibility of long term remodeling of regulatory mechanisms.

When we measured abundance of GFAT1, which catalyzes the rate limiting step of the HBP, we found an increase in its abundance with fasting in CON mice (Figure 3). While this did not occur with a concomitant increase in O-GlcNAcylation levels, it interestingly points to increased faculties to drive O-GlcNAc levels with starvation. In PFKFB2 cKO hearts, there was a state-independent decrease in GFAT1 levels, despite the increase in O-GlcNAc in the fed state. This rules out a role of GFAT1 levels in the observed genotype- and state-dependent changes in O-GlcNAc levels. Additionally, this suggests that O-GlcNAcylation can be upregulated even with considerably decreased GFAT1 levels (Figure 3). This further points to in vivo regulation by coordination between the phosphofructokinase nexus and glucose uptake or availability. Of note, phosphorylated Akt, which is associated with insulin-dependent glucose uptake via GLUT4 translocation to the plasma membrane, is increased in knockout mice but decreases to approximately fed control levels with fasting (Figure 4B). This further supports that glucose uptake (substrate availability) plays an important role in O-GlcNAcylation.

To investigate a potential driver of the counterintuitive decrease in GFAT1 levels in the cKO heart, we measured Sp1 abundance, as this transcription factor not only can be inactivated by O-GlcNAcylation but also regulates GFAT1 expression. The genotype-dependent increase in O-GlcNAc levels supports the potential for a negative feedback loop by which O-GlcNAc downregulates GFAT1 in the cKO mouse. Indeed, we found that O-GlcNAcylation of Sp1 was increased in cKO mice (Figure 3). In addition to regulating GFAT1 levels, Sp1 activates transcription of the gene encoding PDK4^62^. O-GlcNAcylation of Sp1 could thus contribute to downregulation of PDK4 abundance, ultimately decreasing inhibitory phosphorylation of PDH and promoting glucose oxidation in cKO mice^11^. Interestingly, we have also shown that the reciprocal occurs-PDK4 is upregulated in hearts with kinase-dominant PFK-2 overexpression under two different metabolic stressors, fasting and high fat diet feeding^20^. This is of particular interest as hearts from this overexpression model demonstrate decreased UDP-GlcNAc levels under high fat-diet fed conditions^26^. Increased O-GlcNAc also decreases abundance of mitochondrial proteins^27^, as occurs in the PFKFB2 cKO mouse^11^ and impairs mitochondrial function^63^. Sp1 likely influences these changes in mitochondrial protein abundance. The Wende and Abel labs have previously shown that with increased glucose utilization, levels of the Complex I subunit NDUFA9 decreased in an O-GlcNAc and Sp1-dependent manner^30^. This decrease in NDUFA9 abundance also occurs at the protein level in the PFKFB2 cKO model^11^.

Given the established upregulation of O-GlcNAc levels with stress of various forms^47,64,65^, we became interested in how O-GlcNAc levels responded to a 20 minute stimulant stress in the presence and absence of PFKFB2. Strong increases in O-GlcNAc were observed in response to stress across groups. However, in the context of the stimulant stress, fasting did not rescue the genotype-dependent increase in O-GlcNAc in cKO mice (Figure 5J). A plausible explanation for this is a considerable increase in glucose mobilization, uptake, and utilization in the stressed state, irrespective of post-prandial status.

To assess the interplay between cardiac and systemic glucose metabolism, we measured circulating blood glucose and liver glycogen levels in stimulant stressed and unstressed, fed and fasted states. It is established that while metabolically flexible, the healthy adult heart relies predominantly on fatty acid oxidation to meet its energetic demands^66–68^. Previous work performed in the fasted state has even suggested that the heart does not uptake glucose in human patients^69^. Erythrocytes, which lack mitochondria, are obligate glucose users, and the brain also strongly prefers glucose as fuel^70^. Therefore, it is plausible that the heart’s reliance on fatty acid oxidation, especially in the fasted state, is to protect glucose as a substrate for these other tissues. Here we show that in the PFKFB2 cKO model, which upregulates cardiac glucose oxidation, the liver is unable to sustain or upregulate blood glucose levels with acute stress post-fasting. Instead, blood glucose levels decrease significantly (Figure 5B-E). These data highlight the potential maladaptive systemic effects which could arise if the heart were predominantly reliant on glucose.

## Conclusion

Through this study, we set out to test the roles of PFKFB2 loss on mitochondrial metabolic flexibility and to determine the context-dependent role of cardiac PFKFB2 in heart and whole-body glucose homeostasis in response to metabolic and stimulant stress. We also aimed to determine the effects of fed/fasted status, and thus substrate availability, on cardiac O-GlcNAc regulation. We found that PFKFB2 contributes moderately to mitochondrial metabolic flexibility, though the predominant mechanism of PFKFB2-mediated metabolic flexibility is likely due to cytosolic (glycolytic) regulation. In line with changes in glycolytic regulation, PFKFB2 loss drives increased protein O-GlcNAcylation levels in a fed state-dependent manner. We also show that the loss of cardiac PFKFB2 affects systemic glucose metabolism, but not liver glycogen levels, with fasting and stimulant stress. More broadly, from the standpoint of basic scientific research, these data affirm the importance of time of day and post-prandial status in all measurements performed.

## Limitations

In line with our previous work characterizing the cKO model, in which we observed minimal sex-specific differences, we have chosen not to investigate sexes separately in this study. An effort was made to evenly distribute male and female mice between groups.

Isoflurane utilization may impact mitochondrial metabolism. Therefore, respirometry and glucose oxidation studies were performed without isoflurane. All other harvests used isoflurane under the consideration that sedation or light anesthetization would minimize confounding impacts of acute stress on metabolism.

Lastly, we sacrificed fed and fasted mice at two different time points (0700 and 1900 hours). This decision was made because a 12 hour fast over the active period would functionally extend the fast beyond 12 hours, given that the mice were likely fasted during the inactive cycle. Further, this study aims to investigate conditions which recapitulate nocturnal fasting effects. However, it must be considered that several processes, such as PDK4 expression, are likely influenced by a circadian clock component^71^.

## Future Directions

As discussed, we have previously shown that loss of cardiac PFKFB2 promotes structural, electrophysiological, and mechanical remodeling in the heart, as well as a sudden premature mortality. We posited that O-GlcNAcylation is contributing to this electrophysiological dysfunction and mortality as ventricular arrhythmias, Ca2+ handling changes, and sudden death are observed in a transgenic OGT overexpression-mediated model of increased O-GlcNAc^14^. Moreover, O-GlcNAc has been associated with diabetic and hyperglycemia-mediated pro-arrhythmogenic changes in Ca2+ handling ^54^ and repolarization^72^. Therefore, future and ongoing studies in the laboratory will elucidate the fasted state and thus substrate-dependence of PFKFB2 loss on electrophysiological properties in the heart.

## Acknowledgements

We thank Kendra Plafker for guidance on the immunoprecipitation assay.

## Non-Standard Abbreviations

cKO: cardiomyocyte-specific knockout
CON: wildtype control
CPT1: Carnitine palmitoyl transferase 1
FDR: false discovery rate
GC-MS: gas chromatography-mass spectrometry
GFAT: Glutamine fructose-6-phosphate amidotransferase
GLUT4: Glucose transporter type 4
HBP: hexosamine biosynthesis pathway
HOMA-IR: homeostatic model assessment for insulin resistance
LC-MS: liquid chromatography-mass spectrometry
MOPS: 3-(N-morpholino)propanesulfonic acid
O-GlcNAc: O-linked N-acetylglucosamine
OGA: O-GlcNAcase
OGT: O-GlcNAc transferase
PC: palmitoyl carnitine
PCA: principal component analysis
PDH: Pyruvate dehydrogenase
PDK4: Pyruvate dehydrogenase kinase 4
PFKFB2: Phosphofructokinase-2/fructose-2,6-bisphosphatase 2
PFK-1: Phosphofructokinase-1
PFK-2: Phosphofructokinase-2/fructose-2,6-bisphosphatase
PKA: Protein kinase A
Sp1: Specificity protein 1
TAG: triglycerides
TBS: tris-buffered saline
UDP-GlcNAc: uridine diphosphate N-acetylglucosamine

